# Host origin is a determinant of parallel evolution between influenza virus gene segments

**DOI:** 10.1101/2022.02.07.479427

**Authors:** Jennifer E. Jones, Seema S. Lakdawala

## Abstract

Several emerging influenza viruses, including H7N9 and H5N6 viruses, trace their origins to reassortment with H9N2 viruses that contributed internal gene segments. However, the evolutionary constraints governing reassortment of H9N2 viruses remain unknown. In seasonal human influenza A viruses, gene segments evolve in parallel at both the gene and protein levels. Here, we demonstrate that parallel evolution in human H3N2 viruses differs from avian H9 viruses, with both genes and proteins of avian H9 viruses characterized by high phylogenetic divergence. Strikingly, protein trees corresponding to avian H9 polymerase subunits diverge despite known functional constraints on polymerase evolution. Gene divergence was consistent across avian H9 isolates from different continents, suggesting that parallel evolution between H9 gene segments is not dependent on regionally defined lineages. Instead, parallel evolution in H9 viruses was dependent upon host origin. Our study reveals the role of the host in parallel evolution of influenza gene segments and suggests that high reassortment potential in avian species may be a consequence of evolutionary flexibility between gene segments.

## Introduction

Host range is an important factor in viral emergence (1, 2). Influenza A viruses are found in a wide range of hosts, but the natural reservoirs are wild aquatic waterfowl and shorebirds (3, 4). Spillover of avian influenza viruses into humans is rare, but serious zoonotic outbreaks can occur when host range restrictions are overcome (4). In just one example, the H7N9 outbreak on mainland China, a result of reassortment between H7, N9, and H9N2 avian influenza viruses, caused five successive epidemics between 2013 and 2017 with over 1,500 human infections and a mortality rate of 39 percent (5, 6). It is therefore of paramount importance to understand the evolutionary mechanisms that promote emergence of avian influenza viruses in mammalian hosts.

Ecological challenges imposed by the host profoundly impact the spatiotemporal dynamics of viral evolution and emergence (1). Within migratory birds, avian influenza lineages are restricted by host species as well as the migratory routes frequented by these hosts, with little interhost reassortment detected (3, 7). In contrast, in domestic landfowl such as chickens and turkeys, influenza virus subtypes frequently cocirculate and efficiently reassort (8, 9). Adaptation and reassortment of H9N2 viruses in vaccinated farm chickens led to the emergence of the G57 genotype in 2007 with increased antigenicity that overtook all other genotypes by 2013 (10). H9N2 viruses in turn contributed gene segments to other cocirculating influenza viruses, including H7N9, H5N6, and H10N8 viruses (5, 10-13). As of 2016, H9N2 viruses became the dominant subtype in both chickens and ducks in China (14). Therefore, it is critical to consider such factors as geographical restrictions imposed by migratory routes and host species in guiding the evolutionary mechanisms of avian influenza viruses.

Factors intrinsic to viruses are also central to questions of viral evolution and emergence (1). Host range in influenza virus is impacted by several viral properties, including receptor binding specificity and glycosylation of hemagglutinin (HA), stalk length of neuraminidase (NA), and compatibility of the viral ribonucleoprotein complex with host nuclear translocation machinery (15).

Interactions such as those between the viral glycoproteins (HA and NA) or between polymerase subunits (PB2, PB1, PA) functionally constrain influenza virus evolution and may contribute to viral emergence (16, 17). Similarly, packaging signals constrain evolution of the gene segments themselves (18-20). We recently demonstrated that such gene-driven constraints contribute to parallel evolution between gene segments in seasonal human influenza A viruses (21). However, whether evolutionary constraints imposed by interactions between viral genes or proteins extend to the evolution and emergence of avian influenza viruses remains unknown.

We theorized that parallel evolution in influenza viruses was driven by the host. Here, we performed comparative genomics and phylogeography to investigate how the evolution of gene segments differs between human and avian influenza virus strains. We use our established methods to estimate evolutionary convergence between genes and proteins through a proxy tree distance calculation (21). Unexpectedly, we found minimal indication of parallel evolution between gene segments of avian H9 viruses. Instead we provide evidence that the evolution of H9 gene segments converges in a host-specific manner. Our study highlights the role of host origin in shaping influenza virus evolution.

## Results

### Parallel evolution is observed between gene segments of seasonal human H3N2 viruses but not avian H9 viruses

We recently studied whether gene segments of seasonal human influenza A viruses evolve in parallel (21). To achieve this, we examined convergence between gene trees in publicly available human H1N1 and H3N2 virus sequences (**Figure 1A**). We used tree distance as a proxy for convergent evolution between genes, with two gene segments considered converging if the tree distance between them was low. The extent of convergence between pairs of genes varied widely, from robust convergence with tree distances approaching zero to no detectable parallel evolution (21). Parallel evolution between pairs of gene segments differed between viral subtypes but was relatively consistent over time in H3N2 viruses. We therefore examined whether parallel evolution between gene segments of H9 viruses differs from H3N2 viruses found seasonally in human populations.

**Figure 1.**
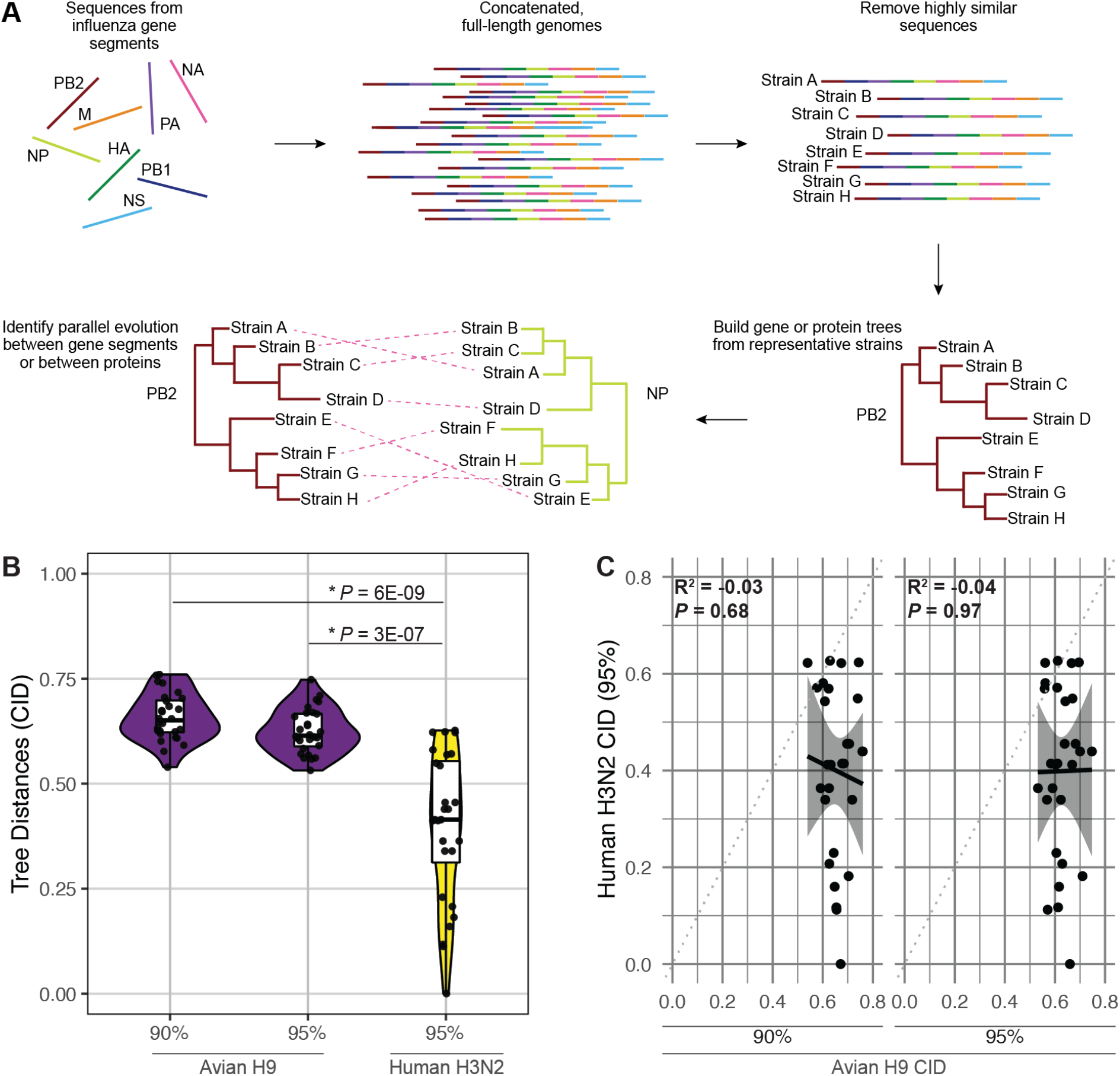
Minimal parallel evolution is found between avian H9 virus gene segments compared to human H3N2 viruses. **A,** Sequences from avian H9 or human H3N2 virus gene segments were obtained from the Influenza Research Database. Gene segment sequences were concatenated into full-length genomes and highly similar sequences were pruned. Maximum likelihood trees of each gene segment were reconstructed from representative strains. Protein trees were reconstructed from coding sequences. Tree similarity was assessed by quantifying the clustering information distance (CID) and used as a proxy for parallel evolution. **B,** CID from avian H9 viruses (90% or 95% similarity thresholds as indicated; purple) compared to human H3N2 viruses (95% similarity threshold; yellow). Each point designates the distance between one pair of gene segment trees. Asterisks (*) indicate *P <* 0.05 (Mann-Whitney *U* test). **C,** Linear regression was performed on pairwise CID. Solid line, best fit. Shaded region, 95% confidence interval. Dotted line, line of identity.

To explore whether parallel evolution between gene segments differs in H3N2 and H9 viruses, we took a similar approach to our previous study (**Figure 1A**), comparing seasonal human H3N2 viruses to avian H9 viruses. In this approach, representative sequences are selected from genomic trees (i.e., species trees) by a sequence similarity threshold (see Methods for additional details). This approach significantly improves gene tree reconstruction by pruning highly similar sequences that don’t resolve well (21). In this study, this approach offered the additional advantage of capturing similar degrees of diversity despite differing sequence availability in human and avian influenza sequences. When a 95% sequence similarity threshold was applied to both human H3N2 and avian H9 viruses, far more sequences remained in avian H9 virus alignments than in human H3N2 virus alignments (200 and 15 respective sequences). Therefore, we examined parallel evolution in avian H9 viruses using trees built from two different sequence similarity thresholds (90% or 95%) to ensure that differences in tree size didn’t artificially inflate differences in tree distance.

Following a modified version of our previously established workflow (21), gene trees were constructed for each of the eight gene segments of each set of viruses (**Figures S1-S2** and **data not shown**). Evolutionary convergence between genes was determined by the clustering information distance (CID) between each pair of gene trees, where CID is inversely correlated to tree similarity and evolutionary convergence. Similar to what we have previously reported (21), tree distances from seasonal human H3N2 viruses ranged from 0 – indicating two identical trees – to 0.63 (**Figure 1B**). In contrast, gene trees derived from avian H9 viruses exhibited significantly higher tree distances than gene trees derived from human H3N2 viruses, ranging from 0.53 to 0.76 (**Figure 1B**). We found no correlation between pairwise CID from avian H9 viruses and human H3N2 viruses (R^2^ = 0.03 to 0.04, **Figure 1C**). These differences were independent of the sequence similarity threshold applied during avian H9 virus gene tree reconstruction (**Figure 1B-1C**). Additionally, tree distances from avian H9 viruses lacked the variation seen in human H3N2 viruses, suggesting little to no preferential evolutionary relationships have formed between individual pairs of gene segments in avian H9 viruses. These data suggest that convergent evolution occurs between gene segments of influenza viruses isolated from human H3N2 viruses, but not avian H9 viruses.

### High divergence is found between avian H9 polymerase subunits

Our initial examination of nucleotide sequences captures evolutionary constraints driven by interactions between protein complexes as well as RNA-RNA interactions. It is surprising that we observed uniformly high divergence between all gene segments in avian H9 viruses, particularly between genes encoding the viral polymerase subunits: PB2, PB1, and PA. Evolution of these three segments is thought to be functionally constrained by protein-protein interactions essential to polymerase function (16, 22, 23). To confirm that parallel evolution does not occur between polymerase subunits in avian H9 viruses, we examined similarity between protein trees. Trees from amino acid sequences of PB2, PB1, and PA from either human H3N2 or avian H9 viruses were constructed (**Figure 2**). As expected, protein trees corresponding to human influenza polymerase subunits were characterized by high convergence, with tree distances ranging from 0.10 to 0.46 (**Figure 2B-C**). In contrast, avian polymerase tree distances were quite high, ranging from 0.76 to 0.78 (**Figure 2A, 2C**). Thus, the polymerase subunits of avian H9 viruses do not exhibit parallel evolution, suggesting that greater flexibility in this complex may be tolerated in avian hosts.

**Figure 2.**
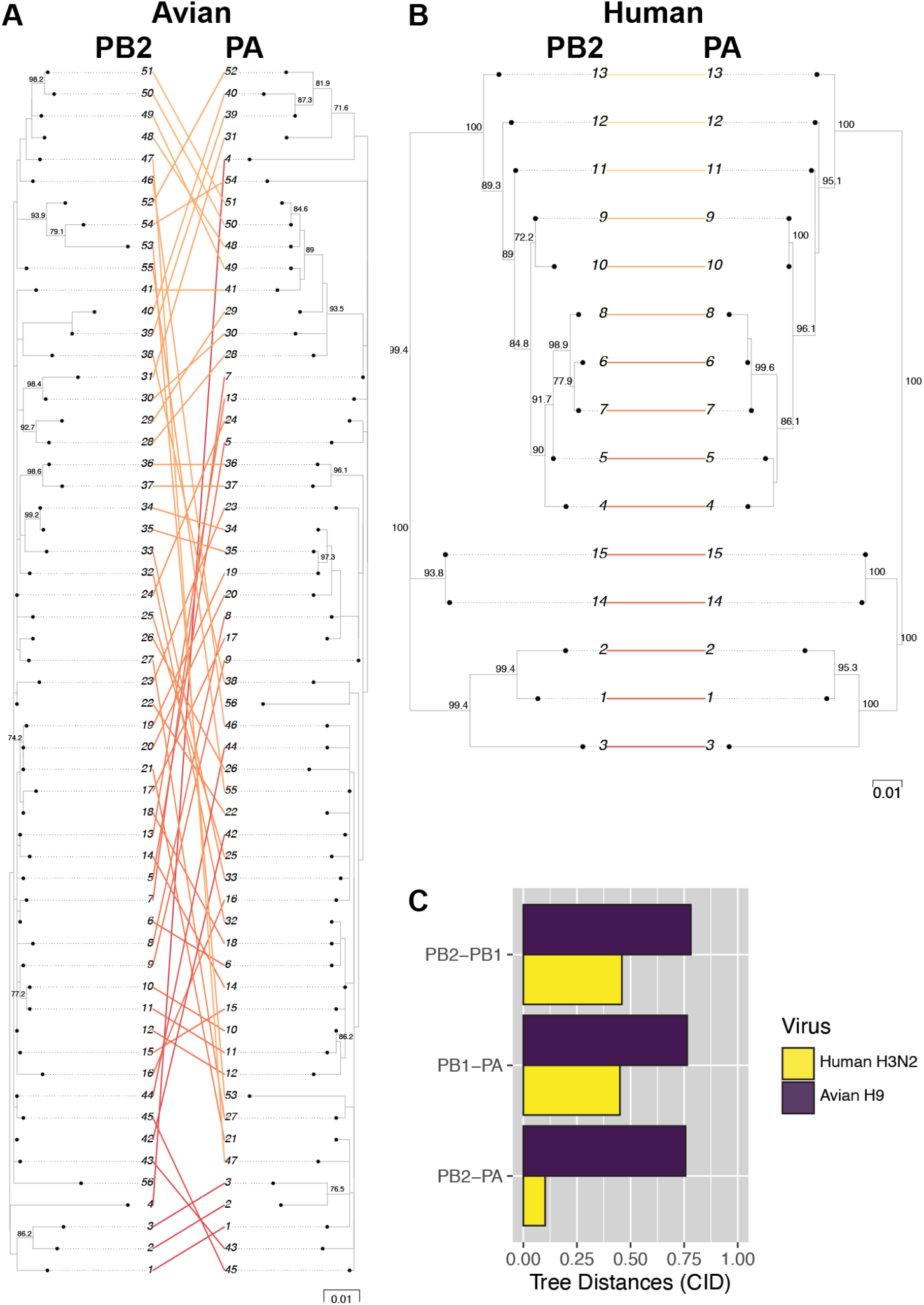
Avian H9 polymerase subunits do not exhibit parallel evolution. Coding sequences corresponding to the gene trees of avian H9 viruses and human H3N2 viruses were used to construct protein trees. **A-B,** Tanglegrams visualizing tree similarity were constructed from the PB2 and PA protein trees for avian H9 **(A)** or human H3N2 **(B)** viruses. Strain names are coded by cluster number. **C,** Pairwise CID for all combinations of polymerase protein trees.

### Divergence of H9 gene segments is consistent across geographical regions

Our investigation takes advantage of the breadth of avian influenza virus sequences available in public databases, but such broad analyses are not without disadvantages. Surveillance of avian H9 viruses is much lower than human H3N2 viruses (**Table S1**). Therefore, sampling bias could distort phylogenetic interpretation. Sampling of avian H9 viruses over time in our dataset was inconsistent, with an inordinately high proportion of sequences coming from 2013 (**Figure S3A**). The disproportionately high representation of sequences from this year coincides with the emergence of H7N9 viruses in China, reflecting heightened poultry surveillance efforts (5, 11). However, our strategy to select sequences from clustering mitigates this sampling bias, with 2013 isolates dropping from 25% of the overall dataset to 13% of sequences selected for phylogenetic reconstruction (**Figure S3B**). Therefore, it is unlikely that our results were greatly impacted by inconsistent surveillance over time.

Another important consideration for avian H9 virus evolution is geographical region. One plausible explanation for the apparent lack of convergence between gene trees in H9 viruses is that parallel evolution between gene segments is lineage specific. We previously reported a similar observation in H1N1 viruses isolated before and after the 2009 pandemic (21). Two geographically distinct H9N2 lineages have emerged from North America and Eurasia (8). Therefore, we examined whether parallel evolution between avian H9 virus gene segments is regionally defined. The vast majority of avian H9 viruses were isolated from Asia (85%), primarily China (**Figure 3A**). Of the remainder, roughly 9% of sequences were isolated from North America, 3% from Africa, 2% from Europe, and less than 1% from South America. We subset avian H9 viruses by continent of origin, excluding viruses from continents with fewer than ten clusters (see Methods for additional details and **Figures S4-S6** for representative trees). When avian H9 viruses were subdivided in this manner, no regional patterns in parallel evolution were detected (**Figure 3B**). However, tree distances were in fact significantly higher in some regions compared to the global dataset. This observation suggests that while lineage-specific differences in tree distances exist, minimal evidence of parallel evolution between gene segments is found in avian H9 viruses from any geographical location.

**Figure 3.**
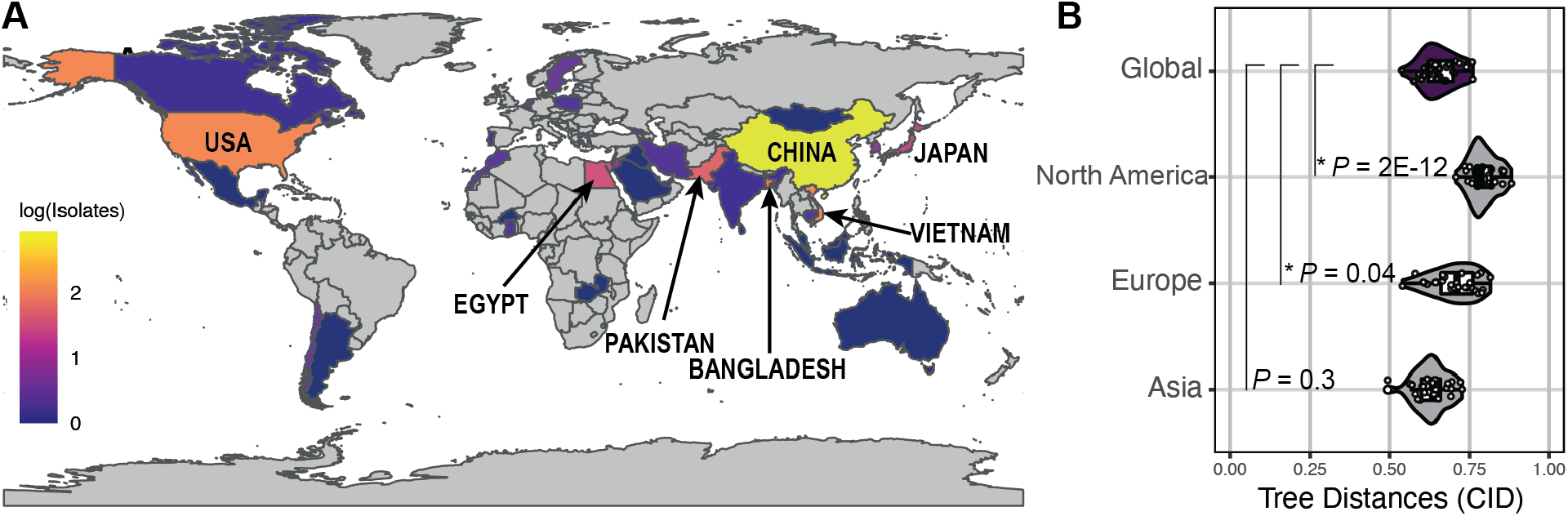
Avian H9 gene tree divergence is independent of region. **A,** Global prevalence of avian H9 viruses. The total number of full-length sequences were log-transformed for visualization. Highest prevalence: China, 861; Vietnam, 120; USA, 118; Bangladesh, 72; Pakistan, 51; Egypt, 34; Japan, 25. **B,** Avian H9 viruses were subset by continent and representative sequences were chosen from each dataset. Maximum likelihood trees of each gene segment were reconstructed from representative sequences. Continents with fewer than ten clusters were excluded. Tree similarity was assessed as described in Figure 1A. Each point designates the distance between one pair of gene segment trees. Asterisks (*) indicate *P <* 0.05 (Mann-Whitney *U* test with Benjamini-Hochberg post-hoc correction).

### Parallel evolution between H9 virus gene segments is dependent upon host origin

Given that geographical region didn’t contribute to evolutionary convergence between gene segments in avian H9 viruses, we examined whether host origin impacts parallel evolution. Host origin could play a role in the overall differences we observed in parallel evolution between gene segments in human H3N2 and avian H9 viruses (**Figure 1B-1C**). Additionally, previous studies suggest that reassortment in wild birds is restricted by host species (7). Therefore, we examined parallel evolution between gene segments of H9 viruses isolated from different hosts, including humans, landfowl, and aquatic birds. Aquatic birds are the natural reservoir of influenza viruses (4), but a sizeable majority of H9 viruses are also found in landfowl such as chickens, turkeys, and quail (**Figure 4A**). Given that tree distances differed in H9 viruses isolated from different continents (**Figure 3B**), we focused our analysis on H9 viruses isolated from Asia, where H9 sequences from humans, landfowl, aquatic birds were all available. Tree distances from human H9 viruses were significantly lower than those obtained from H9 viruses from either set of avian hosts (**Figure 4B** and **Figures S7-S9**). In addition, only tree distances from H9 viruses isolated from humans exhibited the wide range seen in human H3N2 viruses, suggesting evolutionary convergence between gene segments is host dependent. We used linear regression to examine the degree of similarity between individual pairs of gene segments in H9 viruses from different hosts. To our surprise, tree distances from H9 viruses from landfowl were more robustly correlated with those from human hosts than with those from aquatic birds (R^2^ = 0.55 vs. 0.25) (**Figure 4C**). In contrast, tree distances from H9 viruses from aquatic birds were not correlated with tree distances from H9 viruses from human hosts (R^2^ = 0.05) (**Figure 4D**), suggesting that parallel evolution between gene segments in human H9 viruses more closely reflects H9 viruses in landfowl than in aquatic birds. Altogether, these data suggest that parallel evolution between gene segments is dependent on host origin.

**Figure 4.**
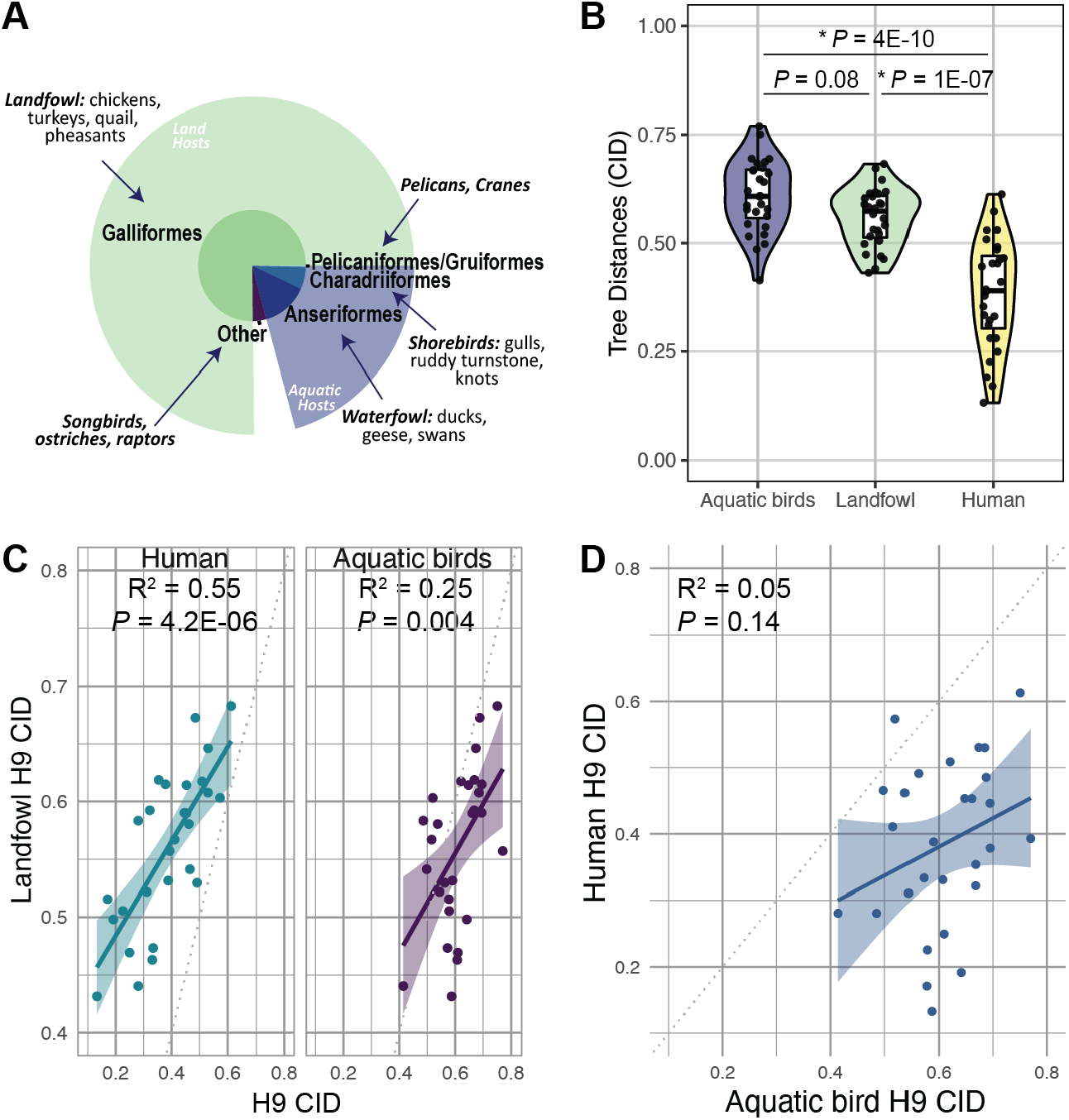
Parallel evolution between gene segments of H9 viruses is dependent upon host origin. **A,** The distribution of H9 viruses across avian taxonomic orders. **B,** CID from Asian-origin H9 viruses isolated from aquatic birds, landfowl, or humans. Asterisks (*) indicate *P <* 0.05 (Mann-Whitney *U* test with Benjamini-Hochberg post-hoc correction).**C-D,** Linear regression was performed on pairwise CID. Solid line, best fit. Shaded region, 95% confidence interval. Dotted line, line of identity. **C,** Comparison of CID in H9 viruses isolated from landfowl to CID in H9 viruses isolated from either humans or aquatic birds. **D,** Comparison of CID in H9 viruses isolated from humans to H9 viruses isolated from aquatic birds.

## Discussion

The mechanistic underpinnings of viral evolution are complex. In the present study, we investigated the interplay between the host niche and functional constraints on influenza A virus evolution. Our results reveal that evolutionary constraints imposed by interactions between viral polymerase subunits in seasonal human influenza viruses do not extend broadly to avian H9 viruses. Instead, the avian H9 polymerase subunits are highly divergent. Surprisingly, the spatial structure of avian influenza lineages established by host migratory routes contributes minimally to parallel evolution of avian influenza gene segments. Instead, gene segment evolution converges in H9 viruses in a host dependent manner. These results highlight the impact of the host niche on influenza virus evolution.

Interactions between viral proteins have long been theorized to impose functional constraints on influenza virus evolution. Coordinated roles between the viral polymerase subunits are well-described and functionally constrain the evolution of each subunit in human influenza viruses (16, 21, 24). Surprisingly, we did not find evidence of parallel evolution between polymerase subunits in avian H9 influenza viruses in this study. Given the success of H9 viruses in the avian population, our data suggest that influenza polymerases may be less evolutionarily constrained in avian hosts than in human hosts. These results are consistent with prior studies demonstrating that the stability of the association between nucleoprotein and the viral polymerase in viral ribonucleoprotein complexes is critical for the evasion of innate immune sensing in human, but not avian, cells (25-27). Overall, these studies may suggest a greater flexibility between viral polymerase subunits in avian hosts that allows for greater success of evolutionarily divergent viruses than in other species.

A lack of parallel evolution between influenza virus gene segments in avian H9 viruses may also have implications for genomic assembly. Selective packaging of all eight influenza gene segments is thought to occur through RNA-RNA interactions between gene segments (18-20, 28). We previously demonstrated that putative intersegmental RNA-RNA interactions could account for some parallel evolution observed between gene segments in human H3N2 viruses (21). Unlike in human H3N2 viruses, gene segments of avian H9 viruses are highly divergent. These results suggest that flexibility may exist in genomic packaging of avian influenza viruses. Evolutionary plasticity in avian influenza gene segments could account for the increased reassortment frequency observed in influenza viruses in avian hosts. Further investigation may reveal a different mechanism of selective packaging of avian influenza gene segments altogether. The problems posed by the unique ecological niche that domestic landfowl afford to avian influenza viruses are a growing concern. Influenza A viruses cocirculate in domestic landfowl and reassort efficiently in these hosts (8, 9). Moreover, the high incidence of coinfection of domestic landfowl with multiple influenza virus subtypes likely influences host range fitness trade-offs. Similar effects have been reported during coinfection with pepper mild mottle virus (1). Here, we discovered that convergence between gene segments in H9 viruses isolated from landfowl more closely mirrors H9 viruses isolated from human hosts than aquatic birds. Our data could be indicative of a role for gene segment convergence in adaptive niches such as landfowl and humans. However, it may instead be the case that evolutionary convergence between gene segments in the aquatic bird reservoir is species-specific. Improved surveillance of H9 viruses in migratory birds will be necessary to discern between these potential mechanisms.

In conclusion, our study reveals the importance of the host niche in influenza virus evolution. It is clear that properties intrinsic to viruses do not shape parallel evolution between gene segments in isolation, but that the host environment can alter the evolutionary trajectories taken. Further investigation of viral evolution in the context of virus-host coadaptation could reveal mechanistic insights into the factors governing viral evolution and emergence.

## Supplemental Information

**Table S1.**
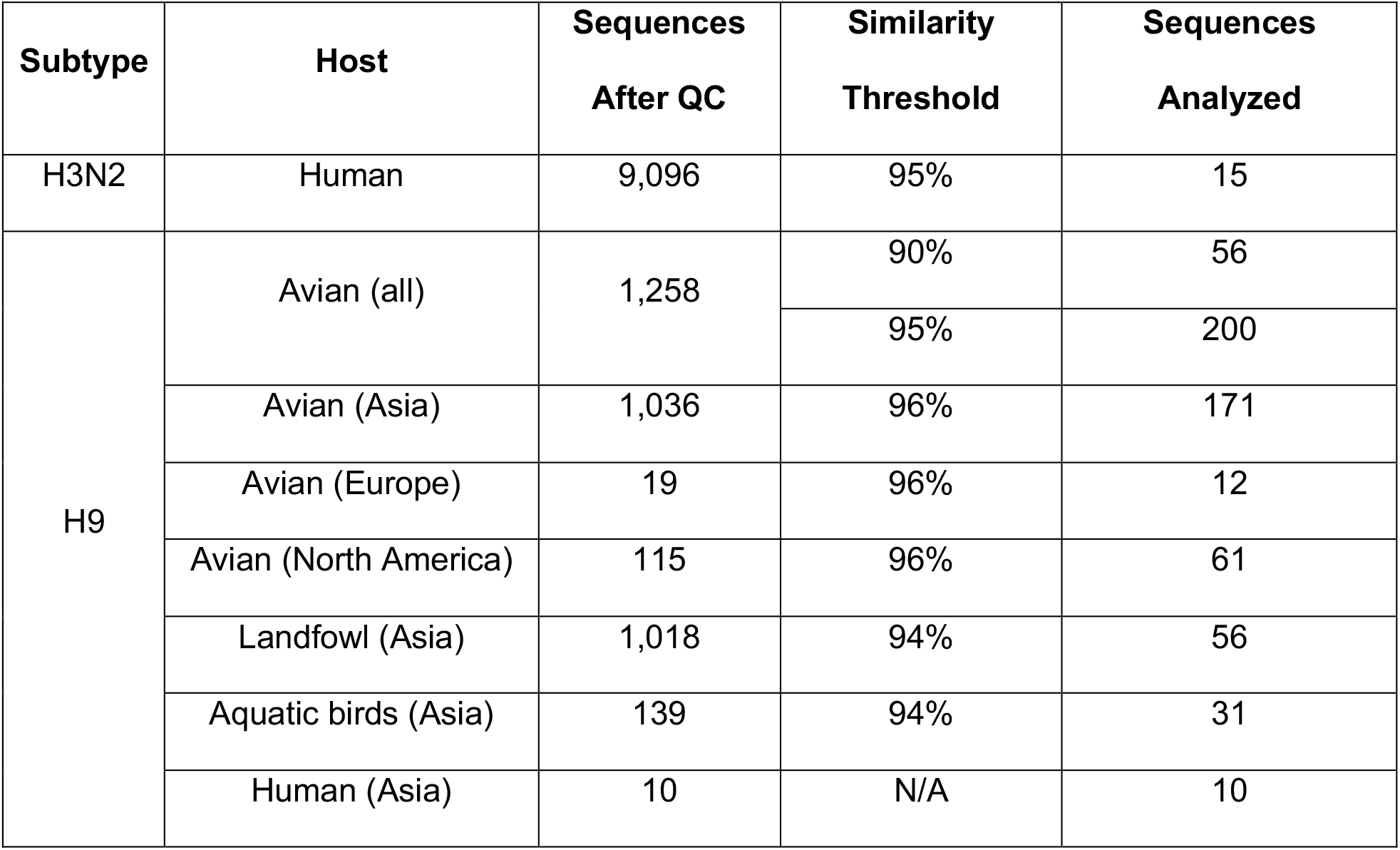
Strains analyzed in this study.

**Figure S1.**
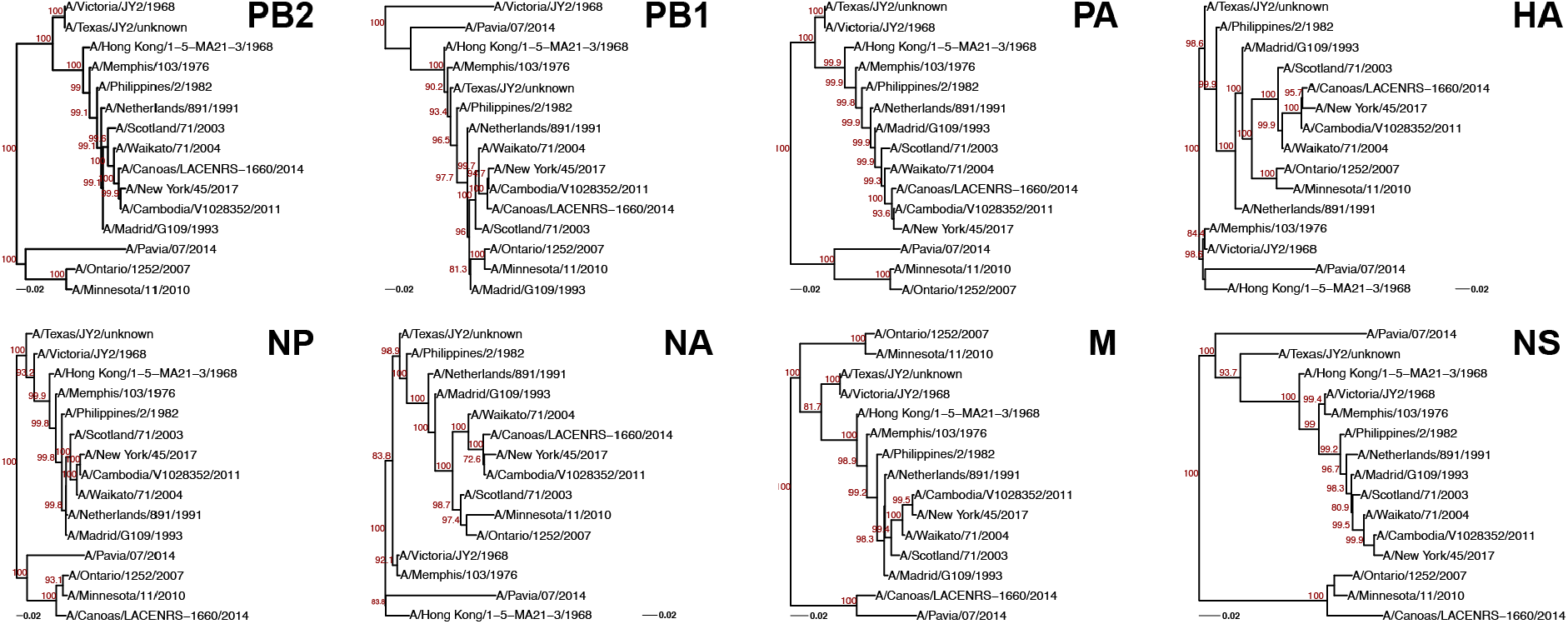
Human H3N2 gene trees. All human H3N2 virus sequences for which full-length genomic sequences were available were downloaded from the Influenza Research Database. A genome tree (i.e., species tree) was constructed from concatenated full-length sequences. Representative sequences were selected by clustering with a sequence identity cutoff of 95%. Maximum-likelihood gene trees were built from these sequences with 1,000 bootstrap replicates. PB2 and NP trees are shown. Bootstrap values greater than 70 are shown in red. Scale bars indicate substitutions per site.

**Figure S2.**
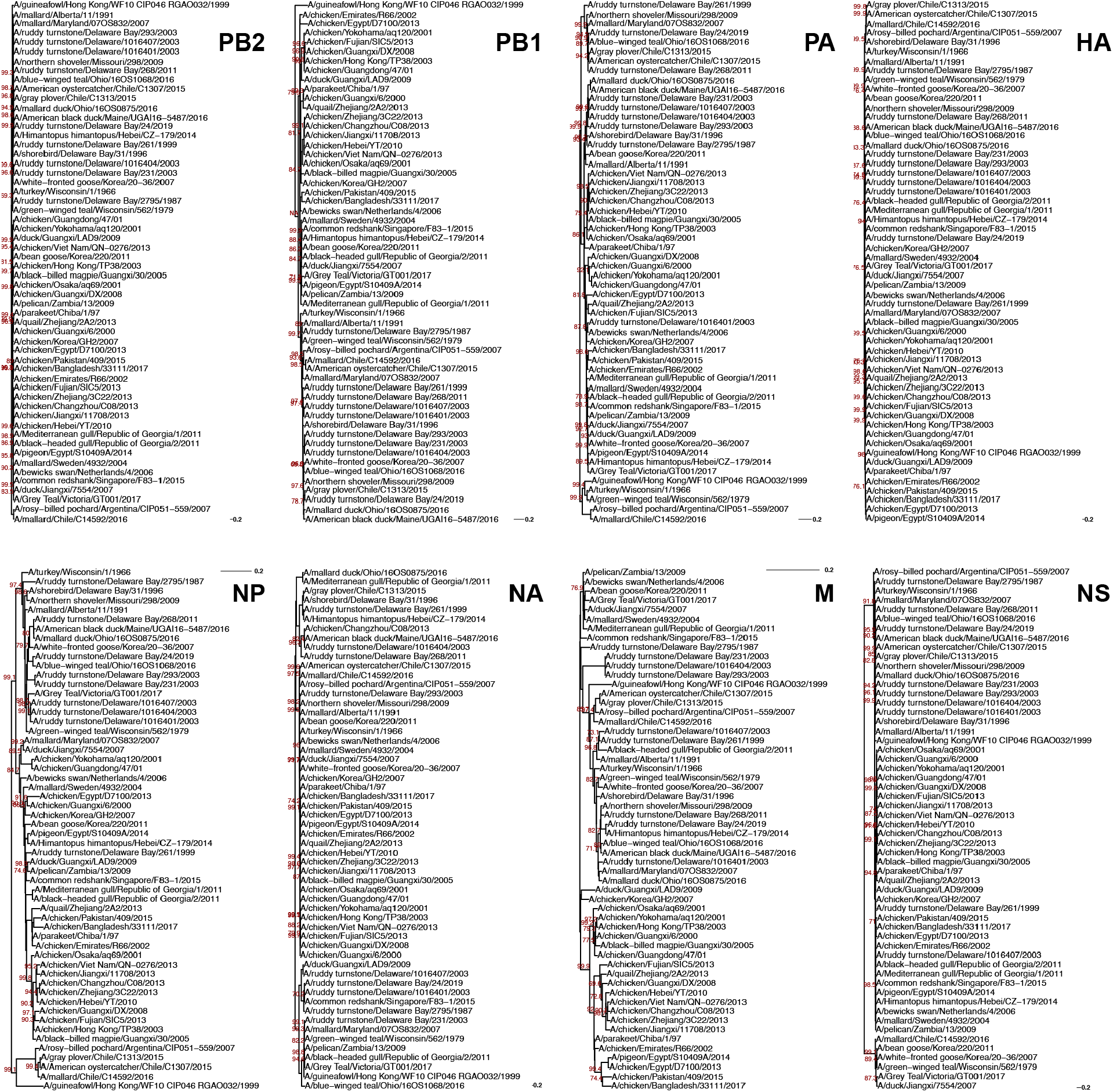
Avian H9 gene trees built with a 90% sequence identity threshold. All avian H9 virus sequences for which full-length genomic sequences were available were downloaded from the Influenza Research Database. A genome tree (i.e., species tree) was constructed from concatenated full-length sequences. Representative sequences were selected by clustering with a sequence identity cutoff of 90%. Maximum-likelihood gene trees were built from these sequences with 1,000 bootstrap replicates. PB2 and NP trees are shown. Bootstrap values greater than 70 are shown in red. Scale bars indicate substitutions per site.

**Figure S3.**
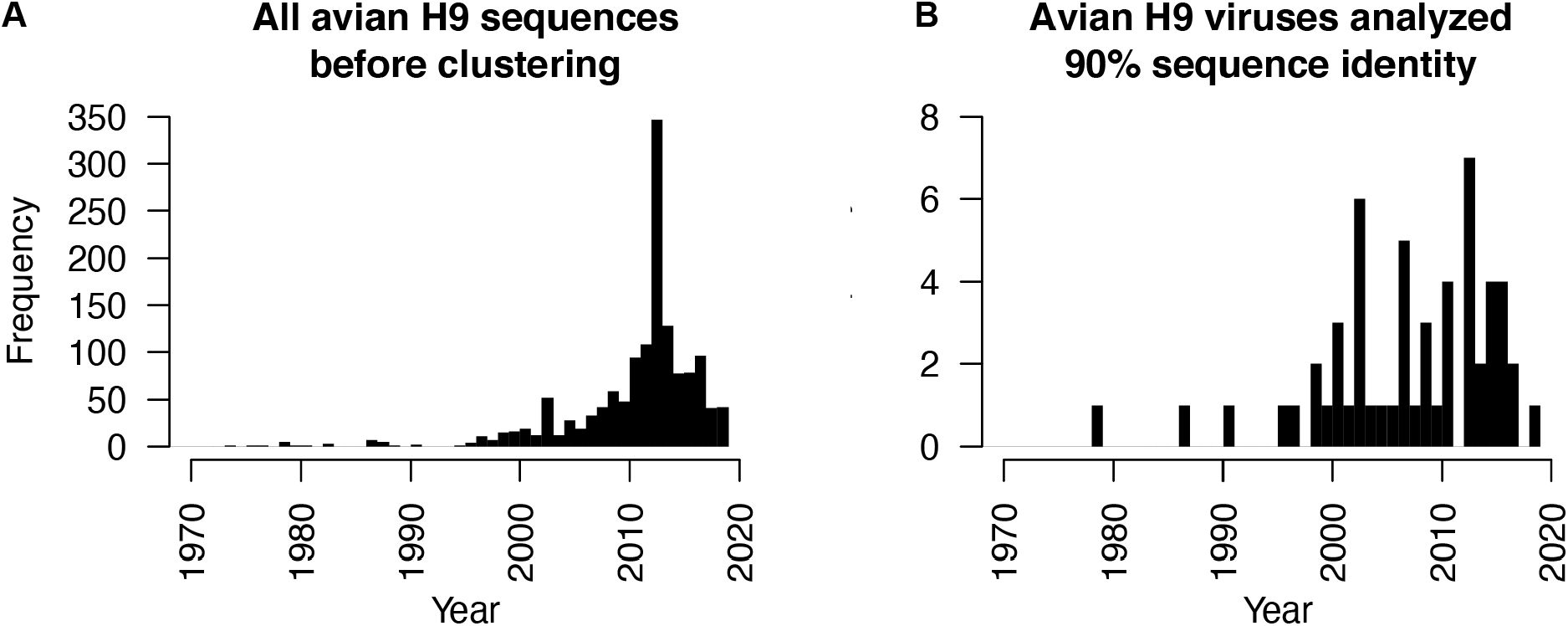
Sampling of avian H9 viruses over time. Frequency of full-length avian H9 virus sequences over time in **A,** all available sequences in the Influenza Research Database, and in **B,** representative sequences selected for construction of gene trees (90% sequence identity shown).

**Figure S4.**
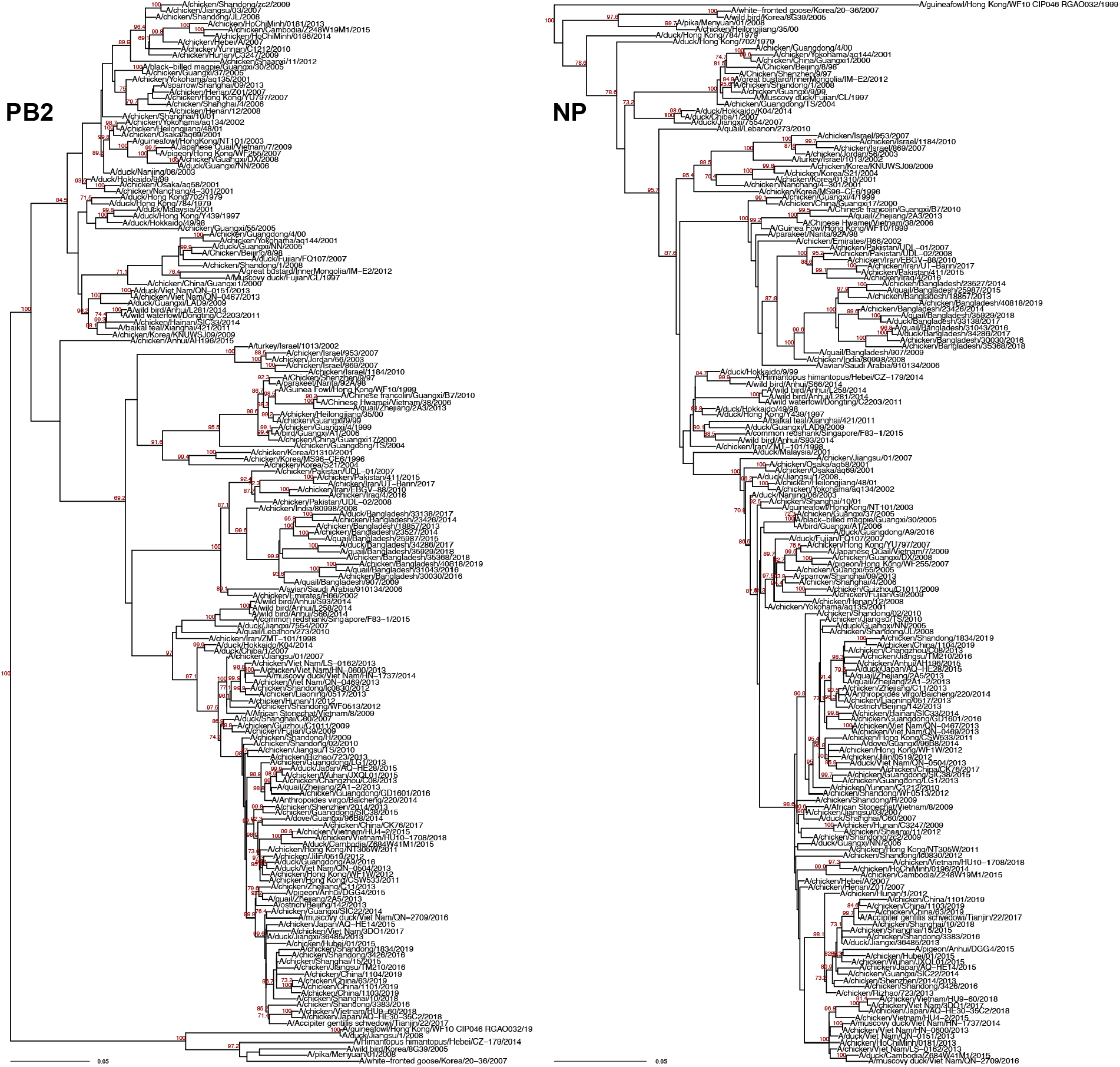
Asian-origin avian H9 gene trees. Avian H9 virus sequences isolated from Asia were selected from all avian H9 viruses described in Figure S2. Representative sequences were selected by clustering with a sequence identity cutoff of 96%. Maximum-likelihood gene trees were built from these sequences with 1,000 bootstrap replicates. PB2 and NP trees are shown. Bootstrap values greater than 70 are shown in red. Scale bars indicate substitutions per site.

**Figure S5.**
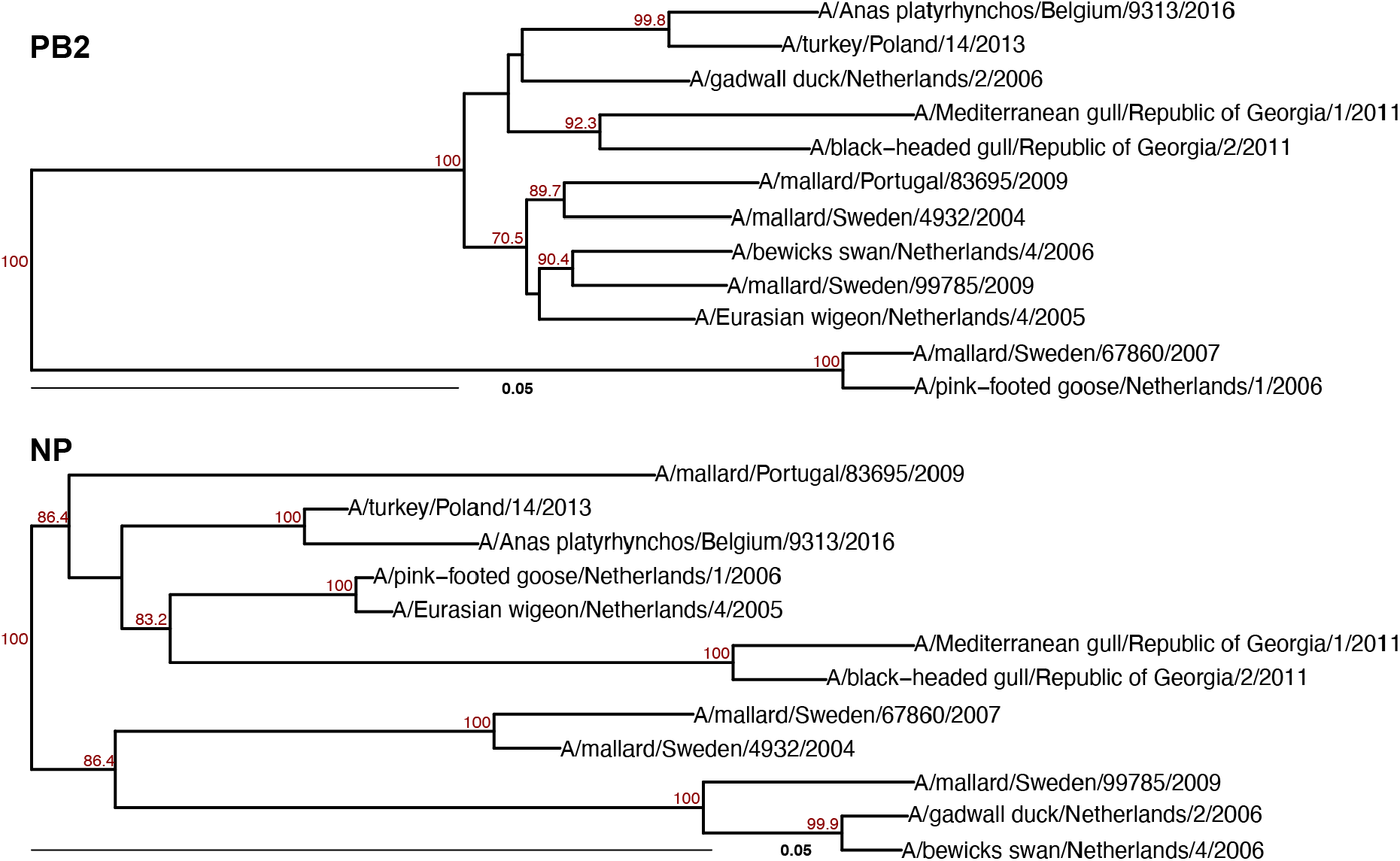
European-origin avian H9 gene trees. Avian H9 virus sequences isolated from Europe were selected from all avian H9 viruses described in Figure S2. Representative sequences were selected by clustering with a sequence identity cutoff of 96%. Maximum-likelihood gene trees were built from these sequences with 1,000 bootstrap replicates. PB2 and NP trees are shown. Bootstrap values greater than 70 are shown in red. Scale bars indicate substitutions per site.

**Figure S6.**
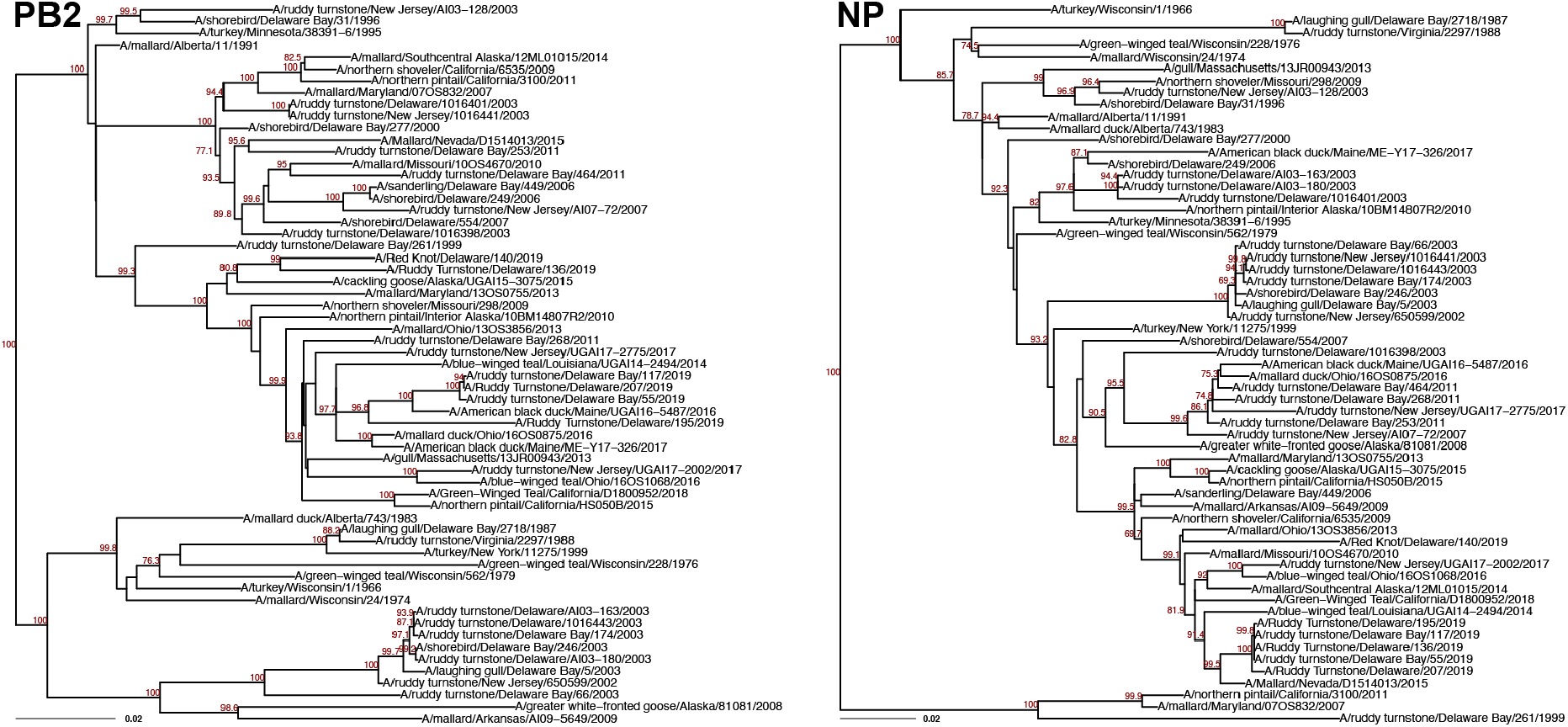
North American-origin avian H9 gene trees. Avian H9 virus sequences isolated from North America were selected from all avian H9 viruses described in Figure S2. Representative sequences were selected by clustering with a sequence identity cutoff of 96%. Maximum-likelihood gene trees were built from these sequences with 1,000 bootstrap replicates. PB2 and NP trees are shown. Bootstrap values greater than 70 are shown in red. Scale bars indicate substitutions per site.

**Figure S7.**
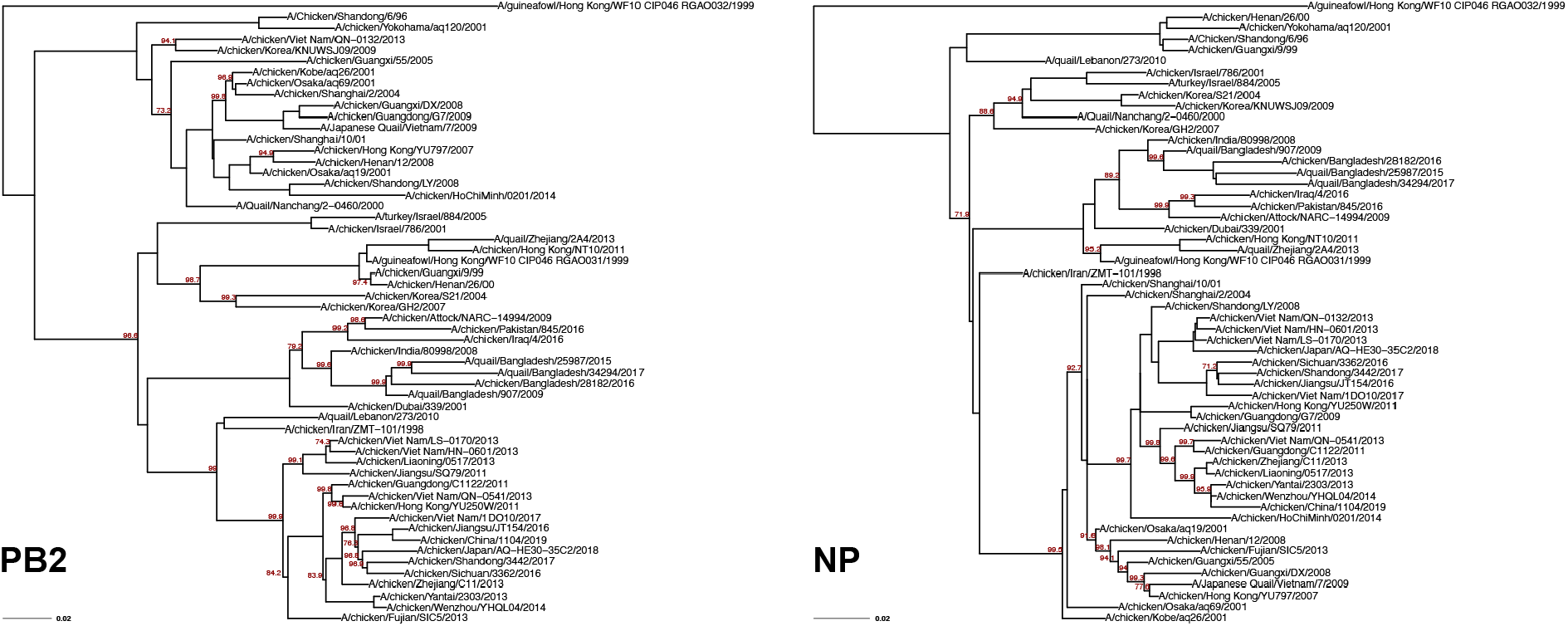
Landfowl-origin H9 gene trees. Landfowl-origin H9 virus sequences isolated from Asia were selected from all avian H9 viruses described in Figure S2. Representative sequences were selected by clustering with a sequence identity cutoff of 94%. Maximum-likelihood gene trees were built from these sequences with 1,000 bootstrap replicates. PB2 and NP trees are shown. Bootstrap values greater than 70 are shown in red. Scale bars indicate substitutions per site.

**Figure S8.**
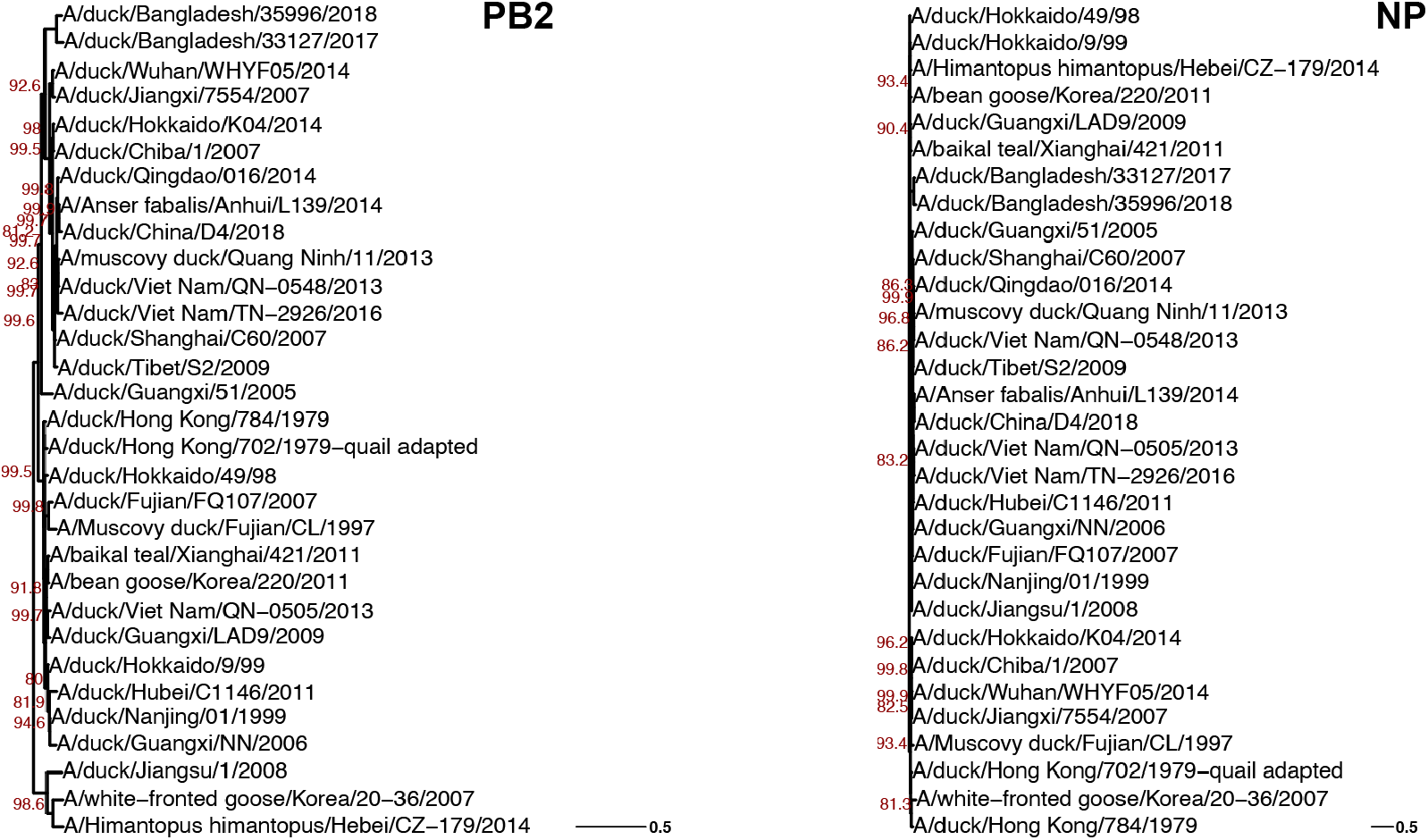
Aquatic bird-origin H9 gene trees. Aquatic bird-origin H9 virus sequences isolated from Asia were selected from all avian H9 viruses described in Figure S2. Representative sequences were selected by clustering with a sequence identity cutoff of 94%. Maximum-likelihood gene trees were built from these sequences with 1,000 bootstrap replicates. PB2 and NP trees are shown. Bootstrap values greater than 70 are shown in red. Scale bars indicate substitutions per site.

**Figure S9.**
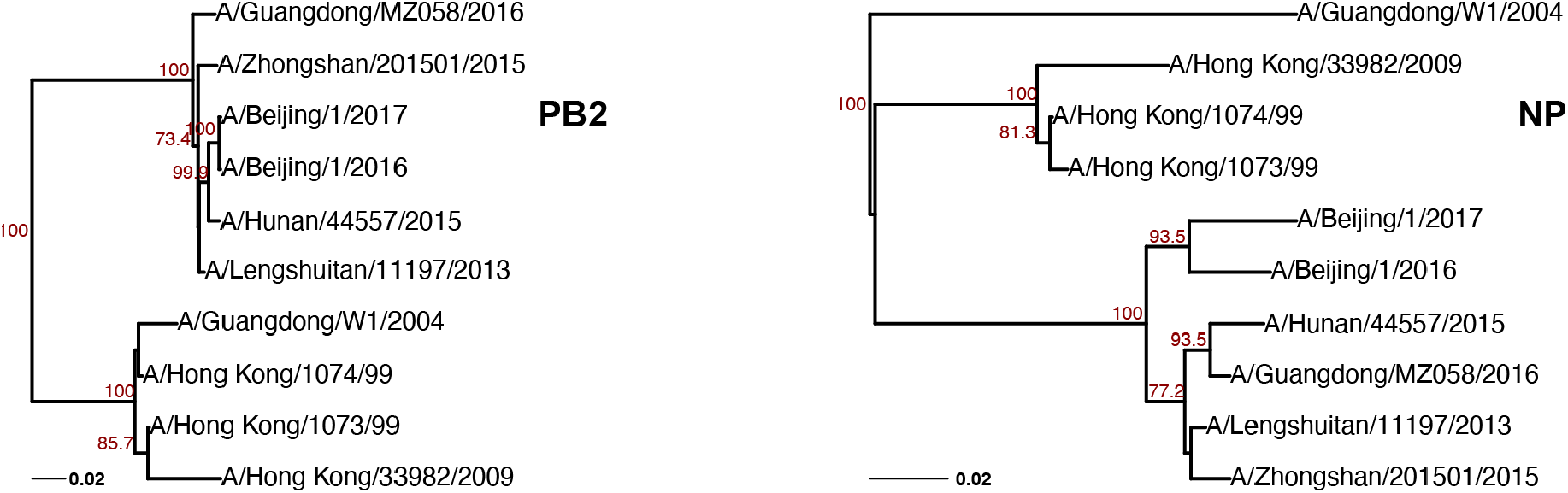
Human-origin H9 gene trees. Human-origin H9 virus sequences isolated from Asia were obtained from the Influenza Research Database. Maximum-likelihood gene trees were built from these sequences with 1,000 bootstrap replicates. PB2 and NP trees are shown. Bootstrap values greater than 70 are shown in red. Scale bars indicate substitutions per site.

## Materials and Methods

### Data Mining and Subsampling

Influenza A virus sequences were mined and sampled as previously described (21). Briefly, FASTA files of genomic segments from avian H9 virus sequences were downloaded from the Influenza Research Database (IRD, http://www.fludb.org) (29) on March 2, 2021. FASTA files of protein coding sequences (CDS) for PB2, PB1, and PA from selected avian H9 virus sequences were downloaded from IRD on June 30, 2021. FASTA files of gene segments from human H9 virus sequences were downloaded from IRD on September 24, 2021. Human H3N2 virus sequences from the dataset described in Jones *et al* were used for comparative genomic analysis of avian H9 and human H3N2 viruses.

Sequences were read into R (version 4.1.0) using the DECIPHER package (version 2.20.0) (30). CDS sequences were translated into amino acid sequences prior to alignment. QC was performed to ensure that sequence duplication, sequencing ambiguity, and incomplete genomes were excluded from analysis. Concatenated alignments comprising all eight gene segments were used to construct species trees for avian H9 and human H3N2 virus sequences by clustering strains into taxonomic units by sequence identity (**Figure 1A**). A sequence identity cutoff of 95% was selected for human H3N2 virus sequences on the basis that this yielded at least ten clusters in the species tree (**Figure S1**). Significant disparities were noted between human H3N2 and avian H9 virus cluster sizes at all cutoffs (e.g., 15 vs. 200 respective clusters in human H3N2 and avian H9 viruses at a cutoff of 95%), so cutoffs of 90% and 95% identity was chosen for avian viruses to ensure that these studies were not biased by tree size. Sampling bias among avian H9 virus sequences was assessed by the year of isolation specified in the FASTA files of sequences after QC and again among sequences used to construct trees (**Figure S3**). Gene and protein trees were built from randomly chosen cluster representatives. Gene trees can be found in **Figures S1-S2**.

### Phylogeography

Phylogeography was performed by analyzing gene trees of avian H9 viruses by continent of origin. All avian H9 strains that remained after QC (1,418 full-length sequences) were assigned to their continent of origin based of the isolation location specified in the FASTA file. Strains from ambiguous locations (e.g., ‘ALB’) were excluded. Sequences were available from all seven continents except Antarctica. These were mapped further to their country of origin and visualized on a world map using the following packages in R: rnaturalearth (version 0.1.0), rnaturalearthdata (version 0.1.0), rgeos (version 0.5-8), and sf (version 1.0-4). Only one sequence was available from Australia and was not analyzed further. Clustering was performed on avian H9 strains from the remaining five continents (Asia, Africa, Europe, North America, South America) as described above, with a sequence identity cutoff of 96% selected for each. South American and African strains each clustered into fewer than ten distinct clusters, so these continents were not analyzed further. Gene trees from the remaining three continents (Asia, Europe, North America) were constructed from randomly chosen cluster representatives (**Figures S4-S6**).

### Host origin

Taxonomical orders represented in avian H9 virus sequences were determined based on hosts specified in FASTA files. Thirteen orders of the Aves class were identified: Galliformes, Anseriformes, Charadriiformes, Pelicaniformes, Gruiformes, Accipitriformes, Passeriformes, Strigiformes, Columbiformes, Falconiformes, Otidiformes, Struthioniformes, Psittaciformes. Sequences with ambiguous or unspecified host species (e.g., ‘Avian’) were excluded. Sequences isolated from Galliformes spp. (including chickens, turkeys, quail, pheasant, guineafowl and Chinese francolin) were designated landfowl-derived (1,065 sequences). Sequences isolated from Anseriformes, Charadriiformes, Pelicaniformes, and Gruiformes spp. were collectively designated aquatic bird-derived (300 sequences). Analysis of landfowl- and aquatic bird-derived H9 virus sequences was restricted to isolates from the Asian continent. Clustering was performed on landfowl-derived and aquatic bird-derived avian H9 sequences as described above, with a sequence identity cutoff of 94% selected for each. Similar numbers of clusters were found despite the considerable differences in overall strain numbers in each group (18 and 33 clusters for landfowl-derived and aquatic bird-derived species trees, respectively). Based on the low number of sequences available for human H9 viruses, all available sequences were used in the reconstruction of gene trees. Representative gene trees are shown in **Figures S7-S9**.

### Tree Reconstruction

Maximum likelihood gene trees were built assuming a general time reversible model of nucleotide substitution using the ape (version 5.5) and phangorn (version 2.7.1) packages. Maximum-likelihood protein trees were built assuming the HIV between-patient model (avian and human PB2 trees, human PA tree) or the FLU model (avian and human PB1 trees, avian PA tree) of amino acid substitution. Best-fit models were approximated by model testing using the AIC criteria. Where indicated by the best-fit model, rates were assumed to vary according to the proportion of invariant sites and/or the discrete Gamma distribution with four rate categories. All trees were assessed for bootstrap support using 1,000 replicates.

### Analysis of Tree Similarity

Tanglegrams, or back-to-back trees matching tips of two trees, were built from pairs of trees using the phytools package (version 0.7-80). The Clustering Information Distance (CID) was calculated with the TreeDist package (version 2.1.1) (31). Statistical significance between tree distances was determined by Mann-Whitney *U* test. Where multiple testing was performed, adjusted *P* values are reported after Benjamini-Hochberg post-hoc correction.

## Code availability

Revised code for analysis of parallel evolution in concatenated, full-length genomic influenza virus sequences is available on GitHub (https://github.com/Lakdawala-Lab/Host-Origin-and-Parallel-Evolution/).

## Acknowledgments

JEJ is supported by a T32 (T32 AI049820) and the Catalyst Award (University of Pittsburgh Center for Evolutionary Biology and Medicine). This work is funded by the National Institutes of Health NIAID (R01 AI139063) and in part with Federal funds from the National Institute of Allergy and Infectious Diseases, National Institutes of Health, Department of Health and Human Services, under Contract No. 75N93021C00015. We thank members of the Lakdawala lab for constructive feedback on this manuscript.

